# Impact of cellular prion protein expression on disease progression and pathology in two mouse models of Alzheimer’s disease

**DOI:** 10.1101/2022.09.25.508690

**Authors:** Silvia A. Purro, Michael Farmer, Elizabeth Noble, Claire J. Sarell, Megan Powell, Daniel Chun-Mun Yip, Lauren Giggins, Leila Zakka, David X. Thomas, Mark A. Farrow, Andrew J. Nicoll, Dominic Walsh, John Collinge

## Abstract

The aggregation of amyloid-β (Aβ) monomers increases their neurotoxicity, and these oligomeric species are thought to be central to the pathogenesis of Alzheimer’s disease. Unsurprisingly for such a complex disease, current Alzheimer’s disease mouse models fail to fully mimic the clinical disease in humans. Moreover, results obtained in a given mouse model are not always reproducible in a different model. Cellular prion protein (PrP^C^) is now an established receptor for Aβ oligomers. However, different groups studying the Aβ-PrP^C^ interaction *in vivo* using a variety of mouse models have obtained contradictory results. Here we performed a longitudinal study in two commonly used AD mouse models using a range of biochemical, histological and behavioural techniques and found similar contradictory results and a possible explanation for the discrepancy. We propose that these two mouse models produce Aβ oligomers with different conformations. Therefore, binding to PrP^C^ and the subsequent activation of toxic signalling cascade will occur only when the Aβ oligomer species with appropriate conformation are present. Hence, it is crucial to select the appropriate model producing the appropriate species of Aβ oligomers to study specific aspects of β-amyloidosis and its downstream pathways. Further conformational characterisation of Aβ oligomers and their binding to PrP^C^ is required to better understand Aβ neurotoxicity.

## Introduction

The neuropathological hallmarks of Alzheimer’s disease (AD) consist of neurofibrillary tangles composed of tau protein, and plaques of amyloid-β (Aβ), in the brain. Although plaques can be toxic to nearby dendrites (Koffie et al. 2009), it has been suggested that the main toxic effects are imparted by soluble Aβ. The Aβ peptide can aggregate in many different arrays and these aggregates or oligomers exist in equilibrium with the monomer and fibril forms (Benilova, Karran, and De Strooper 2012). Currently, many possible receptors for Aβ have been described (Jarosz-Griffiths et al. 2015; Purro, Nicoll, and Collinge 2018; Smith et al. 2019), but finding a cure, or alleviating therapies, have proven elusive. Several years ago, using an unbiased screening approach, Strittmatter’s group identified the cellular prion protein (PrP^C^) as a receptor for Aβ oligomers with nanomolar affinity and further demonstrated that its ablation rescued memory impairment, synaptic dysfunction and perinatal death in the APP_swe_-PS1ΔE9 AD mouse model (APP-PS1) (Lauren et al. 2009; Gimbel et al. 2010). Shortly after, it was reported that PrP^C^ deletion in a different AD mouse model, J20 transgenic mice, did not alter any of the parameters measured, and by contrast actually accelerated premature death in these animals, questioning the role of PrP^C^ in AD (Cisse et al. 2011). Both mouse models overexpress human amyloid precursor protein (APP), albeit under different promoters and harbouring different mutations. APP-PS1 mice express two different PrP^C^ promoter driven transgenes expressing APP with the Swedish mutation (Mullan et al. 1992) together with presenilin 1 (PS1) with an exon 9 deletion (Jankowsky et al. 2004; Jankowsky et al. 2002); whereas J20 mice express APP with Swedish and Indiana (Murrell et al. 1991) familial AD mutations under the platelet-derived growth factor subunit β (PDGF-β) promoter (Mucke et al. 2000). In the subsequent years, many labs (including ours) additionally described two Aβ binding sites present on PrP^C^, identified signalling pathways activated by the interaction, and demonstrated that PrP^C^ ablation, or inhibition mediated by anti-PrP^C^ antibodies, prevents Aβ associated synaptotoxicity *in vitro* and *in vivo*, providing therapeutic proof of principle (Freir et al. 2011; Klyubin et al. 2014; Nicoll et al. 2013; Um et al. 2013; Um et al. 2012; Hu et al. 2014; Corbett et al. 2020). PrP^C^ is now a recognised and validated receptor for Aβ in the AD field.

AD mouse models have been widely used to mostly study either Aβ amyloidosis or tau pathology. Unfortunately, such models do not fully recapitulate neurodegeneration phenotypes. This has been a huge drawback in the field for years. The first generation of AD mouse models overexpress proteins such as mutated APP and/or PS1 to accelerate AD phenotypes within the lifespan of the mouse. Such models differ in several ways, including Aβ plaque burden, localisation, and deposition timing possible due to Aβ kinetics and level of expression, leading to differences in the onset of memory impairments, synaptic dysfunction, neuronal death and presence of tau tangles (Jankowsky and Zheng 2017). Therefore, different pathological AD pathways may be variably present and active in different AD mouse models, and the success of targeting a specific pathway may depend on the characteristics of the individual model. Here, we directly compared the APP-PS1 and J20 mouse lines, studied previously by different laboratories with conflicting results, to independently assess the impact of PrP^C^ deletion on their respective phenotypes, via immunohistochemical, biochemical and behavioural analyses. We found that APP-PS1 mice produce high levels of Aβ oligomers with a conformation that binds to PrP^C^, in contrast with J20 mice, which produce lower levels of PrP^C^-binding oligomers. It is perhaps therefore unsurprising that the J20 AD phenotype is unaffected by PrP^C^ expression. This result suggests that the J20 line is not suitable for investigating the Aβ-PrP^C^ interaction. Moreover, it further highlights the diversity of Aβ oligomers and the necessity to study aggregate conformations, their respective ability to bind to cellular receptors, and their possible function and toxicity.

## Materials and Methods

### Reagents

All chemicals and reagents were purchased from Sigma-Aldrich unless otherwise noted. Synthetic Aβ_1–42_ was synthesized and purified by Dr. James I. Elliott at the ERI Amyloid laboratory Oxford, CT, USA. Peptide mass and purity (>99%) were confirmed by reversed-phase HPLC and electrospray/ion trap mass spectrometry.

### Mice

Work with animals was performed under licence granted by the UK Home Office (PPL 70/9022) and conformed to University College London institutional and ARRIVE guidelines. APP_swe_-PS1ΔE9 mice (APP-PS1, JAX MMRRC Stock# 034829; (Jankowsky et al. 2004)) were obtained from Professor Strittmatter’s laboratory and J20 mice (Mucke et al. 2000) were sourced from The Jackson Laboratory (JAX MMRRC Stock # 034836). Both lines were crossed with either C57BL/6J (Charles River, Margate, UK) or PrP^C^ null backcrossed onto a C57BL/6 background (B6.129S7-Prnp^tm1Cwe^/Orl, EMMA Stock # 01723; (Bueler et al. 1992)) to generate the mouse lines required. Non-transgenic littermates from these crosses were used to populate control groups. Mice were aged and culled at 3, 6 and 12 months old. Groups of 8 male mice were used for biochemical and histological analysis. Mice were anesthetized with isoflurane/O_2_ and decapitated. Brains were removed, divided by a sagittal cut with half brain frozen and the other half fixed in 10% buffered formal saline. Subsequent immunohistochemical investigations were performed blind to sample provenance. For behavioural experiments, 2 cohorts of 15 female and 15 male mice per group were analysed at 2-3, 6-8 and 12-13 months of age. After the last tests, some brains were collected for histological analysis. For Golgi staining, 3 to 5 female brains were collected whole at 12 months of age, see below. The genotype of each mouse was determined by PCR of ear punch DNA and all mice were uniquely identified by sub-cutaneous transponders. RT-PCR was used for determining J20 transgene copy number.

### Immunohistochemistry

Fixed brain was paraffin wax embedded. Serial sections of 5 μm nominal thickness were pre-treated by immersion in 98% formic acid for 8 mins followed by Tris-EDTA buffer for antigen retrieval. All sections were stained with Hematoxylin and Eosin for morphological assessment. Aβ deposition was visualized using 82E1B (cat n.10326, IBL) as the primary antibody, using Ventana Discovery automated immunohistochemical staining machine (ROCHE Burgess Hill, UK) and proprietary solutions. Visualization was accomplished with diaminobenzidine staining.

Histological slides were digitised on a LEICA SCN400F scanner (LEICA Milton Keynes, UK) at ×40 magnification and 65% image compression setting during export. Slides were archived and managed on LEICA Slidepath (LEICA Milton Keynes, UK). For the preparation of light microscopy images, image captures were taken from Slidepath. Publication figures were assembled in Adobe Photoshop.

### Digital image analysis for Aβ quantification

Digital image analysis was performed using Definiens Developer 2.3 (Munich). Initial tissue identification was performed using x10 resolution and stain detection was performed at x20 resolution.

Tissue Detection: Initial segmentation was performed to identify all tissue within the image, separating the sample from background ‘glass’ regions for further analysis. This separation was based on a grey-scale representation of brightness composed of the lowest (darkest) pixel value from the three comprising colour layers (RGB colour model). A dynamic threshold was calculated using the 95^th^ centile which represents the threshold separating the 5% of area with the brightest/highest intensity from the darker 95%; this was then adjusted by −10 (256 colour scale) to ensure accurate tissue separation – this adjustment is necessary to prevent the inclusion of non-tissue regions that, although comprising unstained background, have a reduced pixel value.

Stain Detection: Identification of brown staining is based on the transformation of the RGB colour model to a HSD representation (Der Laak et al. 2000). This provides a raster image of the intensity of each colour of interest (Brown and Blue). Subtraction of the blue stain from the brown stain intensity at each pixel gives a third raster image, Brown+ve, with a positive number where brown stain is prevalent.

All areas with brown staining above 0.15 au, and Brown+ve greater than 0.1, were identified as *Brown Area*. This *Brown Area* was then subdivided to identify *Light Brown Area* < 0.5 au <= *Dark Brown Area*.

Each *Dark Brown Area* was used as a seed for plaques, by growing them into any connected *Light Brown Area*. Plaques were removed if they did not meet several criteria: smaller than 10μm^2^; contained less than 1.5μm^2^ of *Dark Brown Area*; high stain intensity (>0.5 au) and low standard deviation (<0.25 au); area less than 40 μm^2^ with a non-elliptical shape; or area greater than 40 μm^2^ with greater than 70% *dark brown area*.

Tissue Selection: Brain regions were manually selected by hand and plaque and stain coverage data exported per region.

### Golgi staining

Brains were stained using the FD Rapid GolgiStain kit (FD NeuroTechnologies) according the instructions of manufacturer. Stained brains were sliced with a cryostat at 100 μm thickness and developed with solutions provided. After mounting using Permount (TAAB laboratories, Fisher Scientific) at least 15-20 images were taken with a Leica DM2000 LED microscope per animal and spines per dendrite length quantified. Three to five mice per group were analysed, CA1 apical and basal spines were counted using Volocity software (PerkinElmer). Between 1740 and 3960 μm of dendrites were quantified per group.

### Aβ preparations (ADDLs)

Aβ-derived diffusible ligands (ADDLs) were prepared as described previously (Hu et al. 2014). Briefly, 20-25 mg of dry weight peptide was dissolved in 2% w/v anhydrous DMSO for 5 minutes and then diluted to 0.5 mg/ml in phenol red-free Ham’s F12 medium without L-glutamine (Caisson Labs), vortexed for 15 seconds and incubated at room temperature overnight without shaking. After 24-36 h aliquots were tested for the presence of large protofibrillar aggregates using size-exclusion chromatography (GE Healthcare). Fractions containing less than 20% monomer, were centrifuged at 16000 *g* for 20 minutes at 4 °C and the upper 90% of the supernatant collected, snap frozen in liquid nitrogen and stored at − 80°C in aliquots.

### Brain homogenates

Brain samples were homogenised using a cell homogeniser PreCellys24 (Bertin) and whole brain homogenates were prepared at 10% weight in volume (w/v) in PBS with protease and phosphatase inhibitors (Pierce). Bradford quantification of total protein was carried out to ensure similar amounts of proteins were used on the biochemical assays.

### Western blotting

Brain homogenate was thawed on ice for 10 minutes, diluted to a final concentration of 2 mg/ml in PBS, and added to 2x SDS sample buffer. Samples were boiled for 5 minutes then electrophoresed in pre-cast 4-12% NuPAGE Bis-Tris Gels (Invitrogen). Following transfer, nitrocellulose membranes (Amersham GE Lifesciences) were incubated in Licor Odyssey blocking buffer (#927-40000) for 1 h at RT. Membranes were washed 3 × 10 minutes in PBST (PBS, 0.05 % (v/v) Tween-20), then incubated overnight at 4 °C in primary antibodies. APP was labelled using 22C11 Merck Millipore #MAB348 (1:5000), GAPDH was labelled using Sigma G945 (1:50,000) and PrP^C^ labelled using ICSM18 D-Gen (final concentration 3 μg/ml). After incubation in primary antibody, membranes were washed 3 × 10 minutes in PBST, then incubated in secondary antibodies (Odyssey Goat anti-mouse IRDye 800CW or anti-rabbit 680LT) for 1 h at RT. Membranes were then washed twice in PBST, once in PBS and immunoreactive bands were detected and quantified using a Licor Odyssey imaging system (Licor Biosciences).

### Immunoassay to detect PrP^C^ binding Aβ species

To detect PrP^C^ binding Aβ species a plate-based DELFIA (Dissociation-enhanced lanthanide fluorescent immunoassay) was used. HuPrP 23-111 was expressed and purified as described previously for HuPrP 23-231 (Risse et al. 2015). Briefly, inclusion bodies were re-suspended in 6M GdnHCl, β-mercaptoethanol, loaded onto a NiNTA column and refolded by stepwise oxidation. Following elution and dialysis the His tag was cleaved using thrombin, the protein loaded again onto a NiNTA column and eluted in 20mM Bis-Tris, 600mM Imidazole, pH6.5. After dialysis in 20mM Bis-Tris, pH6.5 the protein was stored in aliquots at −80 °C. Thirty microliters of 1 μM human PrP23–111 (10 mM sodium carbonate, pH 9.6) was bound to high binding 384-well white plates (Greiner #G781074) with shaking at 400 RPM for 1 h at 37 °C, washed with 3 × 100 μl of PBST (0.05% Tween-20), blocked with 100 μl Superblock (Thermo Scientific) with shaking at 400 RPM at 37 °C for 1 h and washed with 3 × 100 μl of PBST. Synthetic ADDL preparations were used as standards. Thirty microlitres of benzonase treated 10% brain homogenates (all normalised to the sample with the lowest concentration of protein) were incubated for 1 h at 25 °C with shaking at 400 RPM and washed with 3 × 100 μl of PBST. Aβ oligomers were detected by 30 μl of 0.2 mg/ml 82E1 in DELFIA assay buffer (PerkinElmer) for 1 h at 25 °C with shaking at 400 RPM, washed with 3 × 100 μl of PBST, then incubated for 30 min at 25 °C with shaking at 400 RPM with 9 ng/well of DELFIA Eu-N1 anti-mouse antibody in DELFIA assay buffer (PerkinElmer), washed with 3 × 100 μl of PBST before enhancing with 80 μl of DELFIA Enhancement Solution (PerkinElmer). Plates were scanned for time-resolved fluorescence intensity of the europium probe (λex 320 nm, λem 615 nm) using a PerkinElmer EnVision plate reader.

### Dot blot analysis

One microliter of mouse brain homogenate (2 μg) or synthetic ADDL preparation (5 ng) was spotted directly onto dry nitrocellulose membrane (Amersham) and air dried for 1 h before blocking overnight at 4 °C in 5 % (w/v) non-fat dried milk in PBST (PBS, 0.1 % (v/v) Tween-20). Following three 10 min washes with PBST, membranes were incubated with OC antibody (Millipore, #AB2268) (1:4000 dilution) diluted in 5 % (w/v) non-fat dried milk in PBST overnight at 4 °C. Following three 10 min washes with PBST, membranes were incubated with IRDye 800CW donkey anti-rabbit IgG antibody in Odyssey Blocking buffer (Licor) for 1 h at RT. The membrane was then visualised using an Odyssey scanner. Membranes were subsequently stripped (Restore Plus, Invitrogen) and re-blotted for the loading control β-Actin.

### Multiplex Aβ peptide immunoassay

Levels of Aβ peptides in the mouse brain homogenates were determined using a Multiplex Aβ peptide panel (6E10) immunoassay (Meso Scale Discovery (MSD), Rockville, MD), according to the manufacturer’s instructions. All incubations were carried out at room temperature on a plate shaker at 600 RPM. All standards and samples were diluted in PBS and loaded in duplicate. Before analysis, homogenates were incubated with Guanidine-HCl (6M final concentration) to disaggregate preformed Aβ aggregates. Aβ peptide levels were determined using the MESO QUICKPLEX SQ 120 and analysed using the MSD Workbench 4.0 software.

### Oligomeric Aβ immunoassay

MULTI-ARRAY® 96 well standard bind microplates (MSD) were coated with the monoclonal antibody 1C22 (2 μg/ml) diluted in PBS and incubated at 4 °C overnight. Wells were blocked with 5 % (w/v) Blocker A (MSD) for 1h. Synthetic ADDL preparations were used as standards to generate a twelve point standard curve. All samples and standards were diluted in PBS and loaded in duplicate (25 μl/well). Biotinylated 82E1 (2 μg/ml) diluted in assay diluent (1% Blocker A/PBST) was used for detection. Bound biotinylated 82E1 was measured using SULFO-TAG streptavidin (MSD) diluted in assay diluent. Light emitted from the SULFO-TAG at the electrode surface was detected using the MESO QUICKPLEX SQ 120 imager. All incubations were performed at room temperature on a plate shaker at 600 RPM and all wash steps between incubations were performed using 150 ul PBST, unless stated otherwise. Data were analysed using the MSD Workbench 4.0 software. The limit of detection (LOD) is defined as: LOD = 2.5 x standard deviation of the background. The lower limit of reliable quantification (LLOQ) is defined as the lowest standard with a percentage back interpolation of 100 ± 20 %, a percentage coefficient of variance (CV) ≤ 20 % and a mean blank signal higher than the mean blank signal + (9 * standard deviations of the blank signal). The average of three independent experiments: LLOD 14.1 ± 8.3 (pg/ml) and LLOQ 101.7 ± 55.5 (pg/ml).

### General animal handling

Mice were handled for 10 min a day for several days and habituated to the test environment prior to testing. Only group-housed mice were used. Groups of fifteen to twenty mice were tested sequentially over time for each paradigm, unless otherwise stated. Mice were handled, habituated, trained, and tested at the same time for each experiment.

### Working/recognition memory - Novel Object recognition

This was performed as described (Bozon, Davis, and Laroche 2003). Briefly, mice were tested in a dark cylindrical arena (69 cm diameter) mounted with a MAG300 lamp (DARAY) and Sony Camera Color & F1.3 Lens. 0.7 Lux (Panlabs). Various plastic objects were constructed from interlocking plastic building blocks and were used in all experiments. Pilot studies confirmed all objects were of equal inherent interest. All testing took place in a room with 20 lux lighting and constant background noise.

Animals were habituated to an empty arena for 10 min periods, for 3 consecutive days prior to testing. On day 1 of each experiment (learning phase), two objects were placed in the centre of two 15 cm diameter circles inside the arena. Each mouse was placed in the arena for two periods of 10 min for exploration of the objects with an inter-trial interval of 1 min. One minute later, one of the familiar objects was replaced, with a randomly selected novel one, and retention was tested by placing the mice back in the arena for a 5 min session (test phase). The amount of time spent exploring all objects was measured for each animal with the examiner blind to genotype and time point for the animals. All objects and arena were cleansed thoroughly between trials to ensure the absence of olfactory cues. Criteria for exploration were automated using the Panlabs SMART 3.0 video tracking system where mice had mid-point body mass within a circle of 15 cm around an object. Mice with overt motor symptoms were not used. New sets of objects were used at each time point.

### Spatial Novelty Preference in the Y-Maze

Based on the method by (Sanderson et al. 2011). Spatial novelty preference was assessed in an enclosed Perspex Y-maze with arms of 30 × 8 × 20 cm placed into a room containing a variety of extra maze cues. To promote exploratory behaviour the maze was scattered with a mixture of clean and dirty sawdust (3:1) from the cages of unfamiliar mice of the same sex. Mice were randomly assigned 2 arms (the “start” and the “other” arm) to which they were exposed during the first phase (the exposure phase), for 5 min. Timing of the 5 min period began only once the mouse had left the start arm. The mouse was then removed from the maze and returned to its home cage for a 1 min interval between the exposure and test phases during which the sawdust was mixed and re-distributed around the maze and the divider removed. During the test phase, mice were allowed free access to all 3 arms. Mice were placed at the end of the start arm and allowed to explore all 3 arms for 2 min beginning once they had left the start arm. An entry into an arm was defined using the Panlabs SMART 3.0 video tracking system where mice had mid-point body mass inside an arm. The times that mice spent in each arm were recorded automatically and a novelty preference ratio was calculated for the time spent in arms [novel arm / (novel + other arm)].

### Burrowing – motivational task/innate behaviour

Two hours before the start of the dark period, mice (which had not been food deprived) were placed in individual plastic cages containing a plastic tube 20 cm long × 6.8 cm diameter, filled with 200 g of normal food pellets as described (Cunningham et al. 2003). The weight of pellets remaining in the tube after 24 hr was measured, and the percentage displaced (burrowed) was calculated.

### Statistical Analysis

All statistical analysis and graphs were generated using the statistical package GraphPad PRISM v7 (GraphPad Software, Inc., La Jolla, USA). For multiple comparisons, graphs depict median values and Kruskal-Wallis was used and corrected for multiple comparisons using Dunn’s multiple comparison test, except for behavioural tests where two way ANOVA followed by Sidak’s test was used. Statistical significance was set to P < 0.05.

## Results

### Aβ aggregation is independent of PrP^C^ expression

APP-PS1 mice overexpress APP carrying the Swedish mutation plus PS1 with deletion of exon 9 (Jankowsky et al. 2002; Jankowsky et al. 2004). In contrast, J20 mice overexpress only APP with Swedish and Indiana mutations (Mucke et al. 2000). In order to assess the role of PrP^C^ in both AD mouse models, we crossed the APP-PS1 and J20 mice with a PrP^C^ knock-out (KO) mouse line. Wild-type or PrP^C^ KO littermates generated without expressing APP or PS1 transgenes were used as controls (Figure 1A). PrP^C^ expression did not alter the expression of APP, nor did the overexpression of the mutated genes APP or APP/PS1 induce any changes in PrP^C^ levels (Figure 1B and 1C).

**Figure 1.**
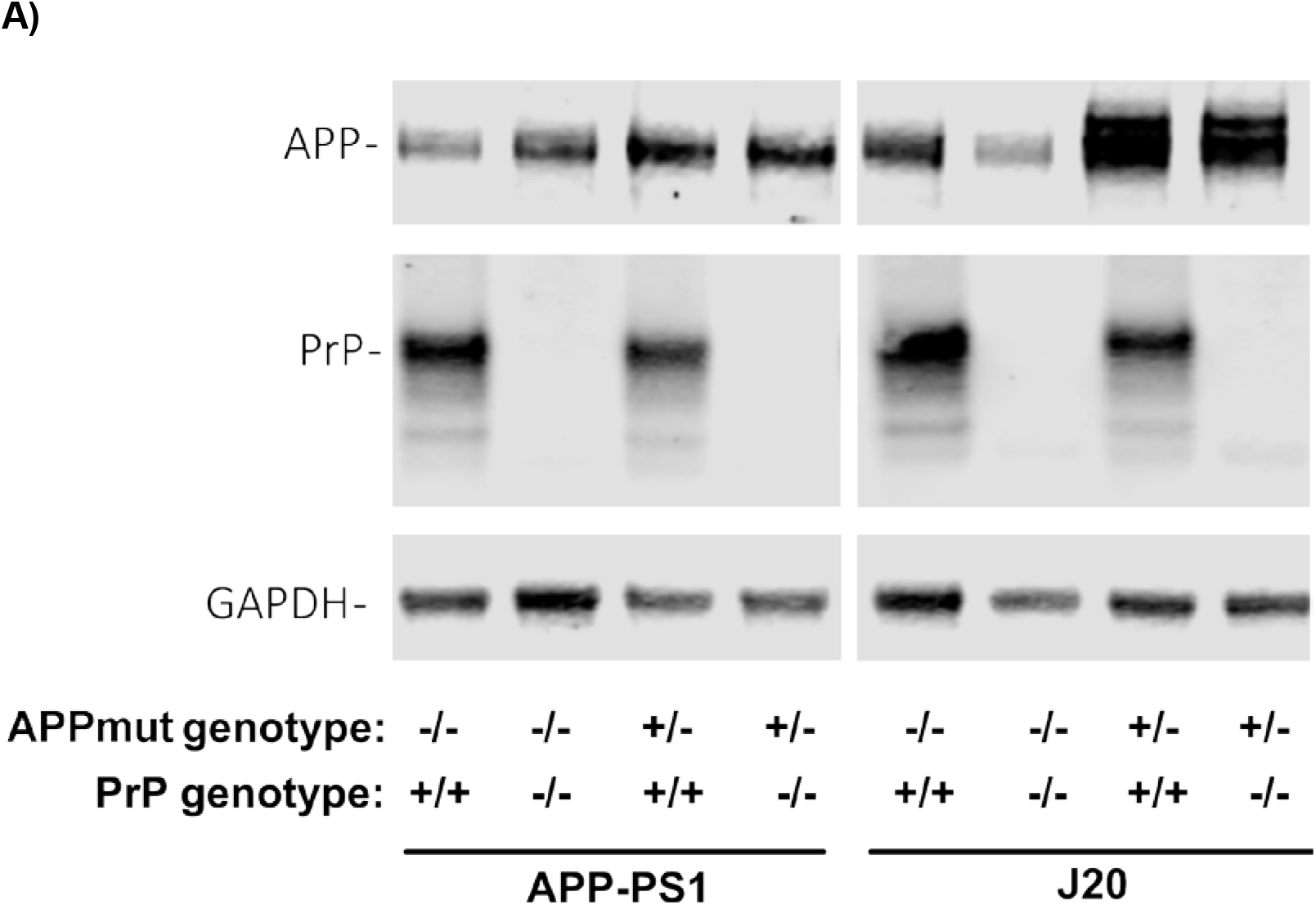
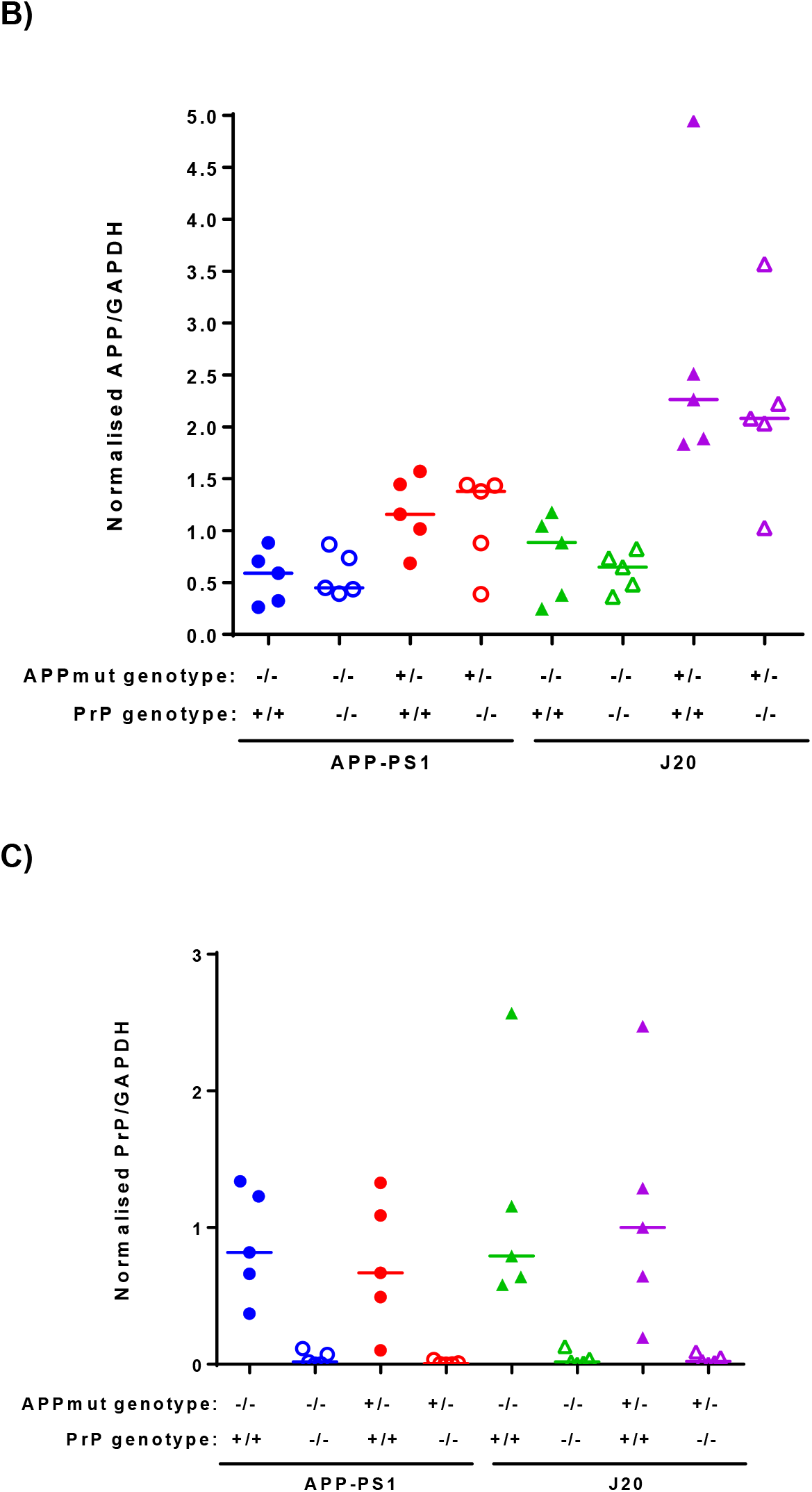
Knockout of PrP^C^ does not affect total APP levels in wild type or AD-model mice. **A)** Total brain homogenate from 12-month old mice across 8 genotypes were analysed via western blot. APP was labelled using the N-terminal monoclonal antibody 22C11, and PrP^C^ labelled using ICSM18. Homogenate from a single sample was loaded on every gel analysed (n = 5) and used to normalise protein quantifications between gels. Values reported are the ratio of APP/GAPDH within each sample. **B)** Quantification of APP expression levels determined by western blot. J20 mice showed significantly higher total APP expression than APP-PS1 mice irrespective of PrP^C^ status (APP-PS1 vs J20, p=0.024), however no significant differences were observed between the PrP^C^ +/+ and -/- variants of any APP genotype. **C)** Quantification of PrP^C^ expression levels determined by western blot. No significant differences in PrP^C^ expression were observed between APP genotypes. Deletion of PrP^C^ resulted in a significant difference on PrP^C^ expression levels for all the lines used in this study, when compared to their respective controls (WT vs PrP^C^ KO, p=0.036; APP-PS1 vs APP-PS1 PrP^C^ KO, p=0.019; WT vs PrP^C^ KO, p=0.019; J20 vs J20 PrP^C^ KO, p=0.026).

For the four mouse lines (APP-PS1, J20, APP-PS1 PrP^C^ KO and J20 PrP^C^ KO) and their respective control littermates (WT and PrP^C^ KO) we collected brain samples at 3, 6 and 12 months of age and, then examined Aβ species and aggregation using immunohistochemical and biochemical techniques.

Deposition of Aβ in the brains of J20 mice is only visible after 6 months of age and plaques are almost fully concentrated in the hippocampus, corpus callosum and cortex. Minimal plaques appear in the cerebellum after 12 months of age. By contrast, plaques in APP-PS1 mice are spread over the cerebellum, olfactory bulb, hippocampus, corpus callosum, cerebral cortex and other areas of the brain. APP-PS1 whole brain sagittal sections exhibit twice as many plaques at 12 months than the J20 mice (APP-PS1 median: 2815 plaques, J20 median: 1315 plaques, p=0.014) whilst brain area plaque coverage is comparable between lines (APP-PS1 median: 1.16 %, J20 median: 0.88 %, p=0.75) (Figure 2A-C). Ablation of PrP^C^ did not alter the number, localisation or area covered by plaques in either AD mouse model (Figure 3A-C), in agreement with previously published results (Gimbel et al. 2010).

**Figure 2.**
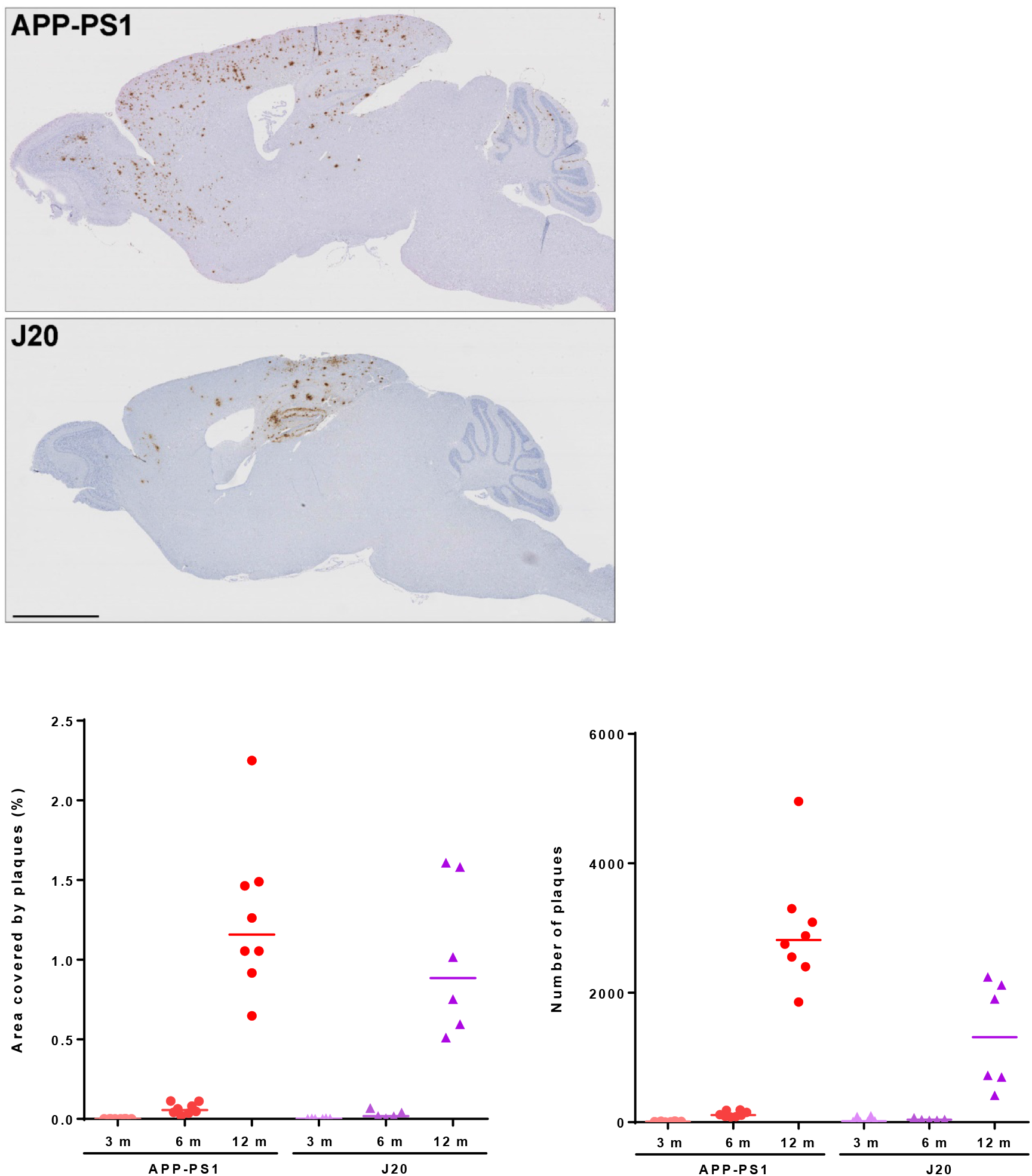
Progressive brain deposition of Aβ plaques in APP-PS1 and J20 mice. **A)** Representative images of APP-PS1 and J20 12-month old mice stained with 82E1b anti-Aβ antibody. **B)** Quantification of Aβ plaques area during aging. **C)** Quantification of Aβ plaques number. APP-PS1 mice exhibited a higher number of plaques at 12 months of age (APP-PS1 vs J20, p=0.014). Scale bar: 2 mm.

**Figure 3.**
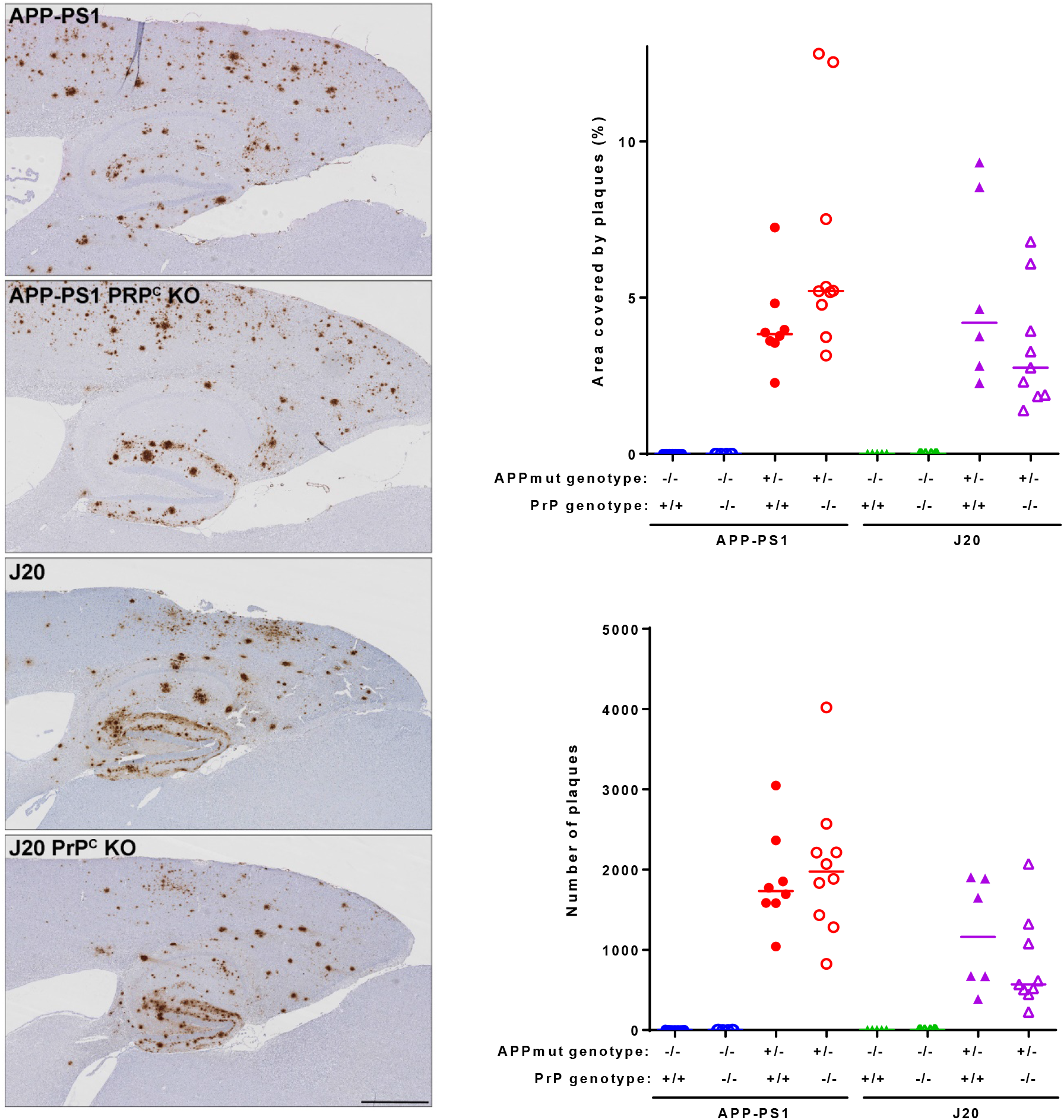
Deposition of Aβ plaques in APP-PS1 and J20 mice is independent of PrP^C^ expression. **A)** Representative images showing hippocampus, cortex and corpus callosum at 12 months of age of the four mouse lines studied. **B)** Quantification of the area covered by plaques on the above mentioned areas. For all the APP mutant lines there was a significant difference in area covered when compared to their respective controls (WT vs APP-PS1, p=0.02; PrP^C^ KO vs APP-PS1 PrP^C^ KO, p=0.0005; WT vs J20, p=0.0006; PrP^C^ KO vs J20 PrP^C^ KO, p=0.037) and, similar amounts when compared APP mutant lines to their respective ablated PrP^C^ line (APP-PS1vs APP-PS1 PrP^C^ KO, p=0.75; J20 vs J20 PrP^C^ KO, p>0.99). **C)** Quantification of the number of plaques on the above mentioned areas. Transgenic mice exhibited a higher number of plaques compared to their respective wild-type littermates (WT vs APP-PS1, p=0.005; PrP^C^ KO vs APP-PS1 PrP^C^ KO, p=0.002; WT vs J20, p=0.0008; PrP^C^ KO vs J20 PrP^C^ KO, p=0.031) and no significant difference when compared to their respective ablated PrP^C^ line (APP-PS1vs APP-PS1 PrP^C^ KO, p>0.99; J20 vs J20 PrP^C^ KO, p>0.99). Scale bar: 700 μm.

Total Aβ peptides in whole brain homogenates collected at 6 and 12 months were quantified by immunoassay. At 12 months old, APP-PS1 mice had significantly more Aβ_42_ than J20 mice (for Aβ_42_ APP-PS1 median: 223 ng/mg, J20 median: 55 ng/mg, p=0.04), but did not reach significance for the Aβ_40_ peptide (for Aβ_40_ APP-PS1 median: 113 ng/mg, J20 median: 16 ng/mg, p=0.14); with PrP^C^ having no impact on Aβ peptide levels in either model (Figure 4). The amount of Aβ_40_ and Aβ_42_ peptides in APP-PS1 mice was almost 95% lower at 6 months versus 12 months of age (data not shown).

**Figure 4.**
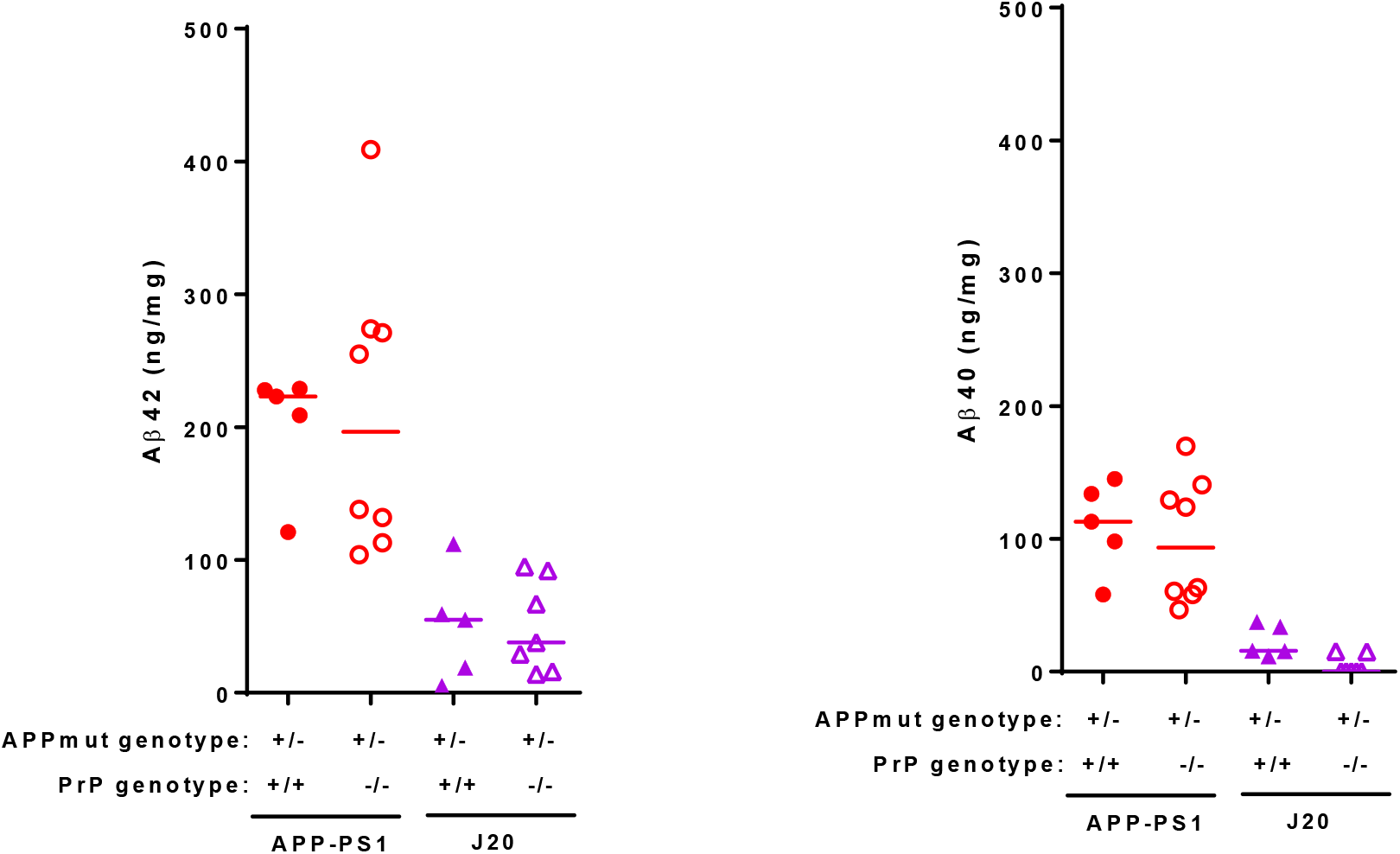
Immunoassays demonstrate that total Aβ peptide levels in APP-PS1 and J20 mouse brain are independent of PrP^C^ expression. Total brain homogenates from mice at 12 months of age were analysed by multiplex Aβ peptide panel (6E10) immunoassay from MSD. **A)** APP-PS1 expressing or not PrP^C^ presented higher levels of Aβ_42_ than J20 samples (APP-PS1 vs J20, p=0.04; APP-PS1 PrP^C^ KO vs J20 PrP^C^ KO, p=0.004). **B)** Quantification of Aβ_4o_ levels revealed no changes due to ablation of PrP^C^ in any of the mouse lines. Levels of Aβ_4o_ were not significantly different between APP-PS1 and J20 mice (APP-PS1 vs J20, p=0.14).

Similarly, when the levels of Aβ oligomers were quantified using the 1C22 immunoassay that does not recognise Aβ monomers, we found that APP-PS1 mice had significantly higher levels than their wild-type littermates, whereas there was no detectable significant difference between the J20 mice and their wild-type littermates, at 12 months of age (WT vs APP-PS1, p=0.035; WT vs J20, p=0.25). PrP^C^ expression had no impact on the levels of 1C22-reactive Aβ oligomers in either APP-PS1 or J20 transgenic lines (Figure 5). Together, these results suggest that distinct Aβ conformers and/or aggregation mechanisms exist in the respective transgenic lines, as the plaque burden at 12 months old is relatively similar, but levels of soluble Aβ differ greatly.

**Figure 5.**
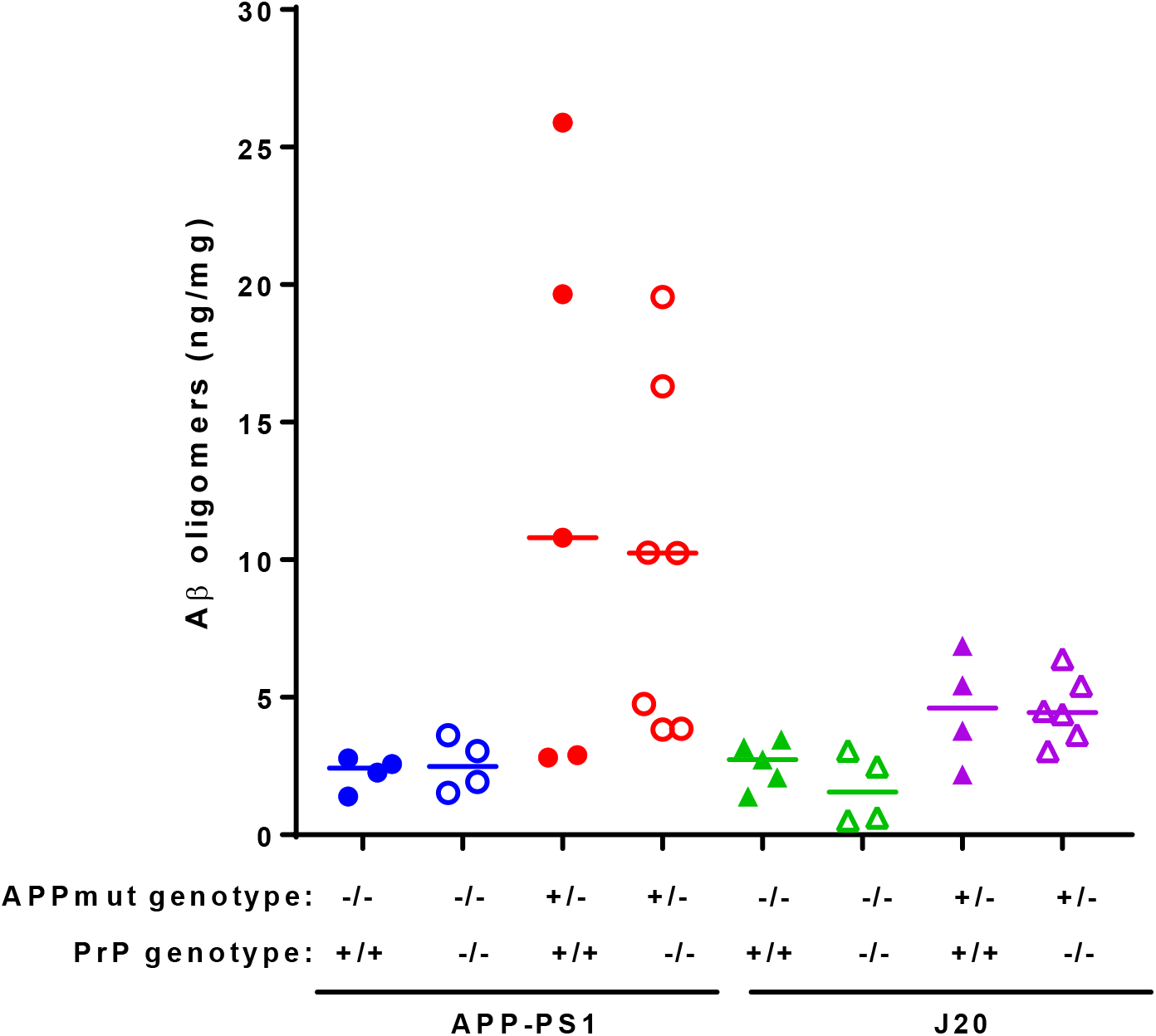
Quantification of Aβ oligomers present in APP-PS1 and J20 mouse brain do not change with PrP^C^ expression. Total brain homogenates were assayed using 1C22 anti-Aβ oligomer antibody (WT vs APP-PS1, p=0.035; APP-PS1 vs APP-PS1 PrP^C^ KO, p>0.99; WT vs J20, p=0.25; J20 vs J20 PrP^C^ KO, p>0.99).

Analysis of the levels of Aβ oligomers capable of binding PrP^C^ revealed no significant differences between J20 mice and their respective wild type controls. In contrast, APP-PS1 samples contained significantly higher levels of these Aβ oligomers, with no differences detected between mice with *Prnp +/+* and *Prnp -/-* backgrounds (Figure 6). We then characterised the conformation of the Aβ oligomers, using the OC antibody which recognises parallel, in register fibrils (distinct from the A11 antibody, which binds to anti-parallel Aβ structures (Kayed et al. 2007; Glabe 2008). Interestingly, only APP-PS1 mice but not J20 mice, presented a significant amount of these OC-Aβ oligomers (WT vs APP-PS1, p=0.012; WT vs J20, p=0.87) (Figure 7). It has been suggested that Aβ oligomers that are able to bind to PrP^C^ have an OC conformation (Nicoll et al. 2013; Madhu and Mukhopadhyay 2020). Levels of OC-Aβ oligomers did not change after ablation of PrP^C^ in either mouse line (Figure 7). Interestingly, only APP-PS1 mice, and not J20 mice, showed a positive and significant correlation between total amount of Aβ oligomers and oligomers that bind to PrP^C^ (Fig 8). This is in agreement with previous studies which showed that J20 mice produce mainly A11-Aβ oligomers (Liu et al. 2015).

**Figure 6.**
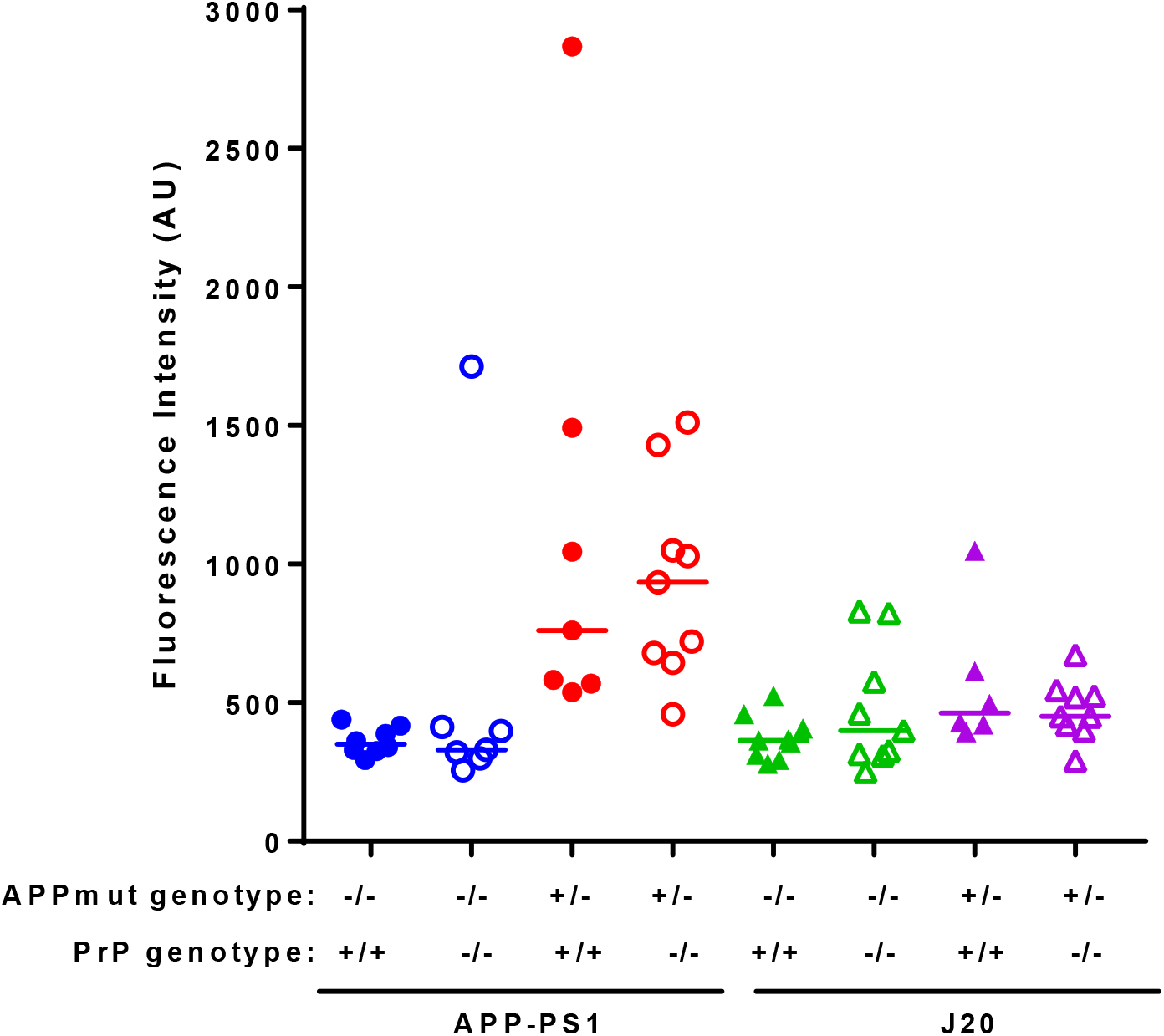
Aβ oligomers that bind to PrP^C^ are present in APP-PS1 at higher levels than in J20 mouse brain, but do not change with PrP^C^ expression. Total brain homogenates were analysed by DELFIA immunoassay to detect PrP^C^-binding Aβ species (APP-PS1 vs J20, p=0.037; WT vs APP-PS1, p=0.009; APP-PS1 vs APP-PS1 PrP^C^ KO, p>0.99; WT vs J20, p=0.12; J20 vs J20 PrP^C^ KO, p>0.99).

**Figure 7.**
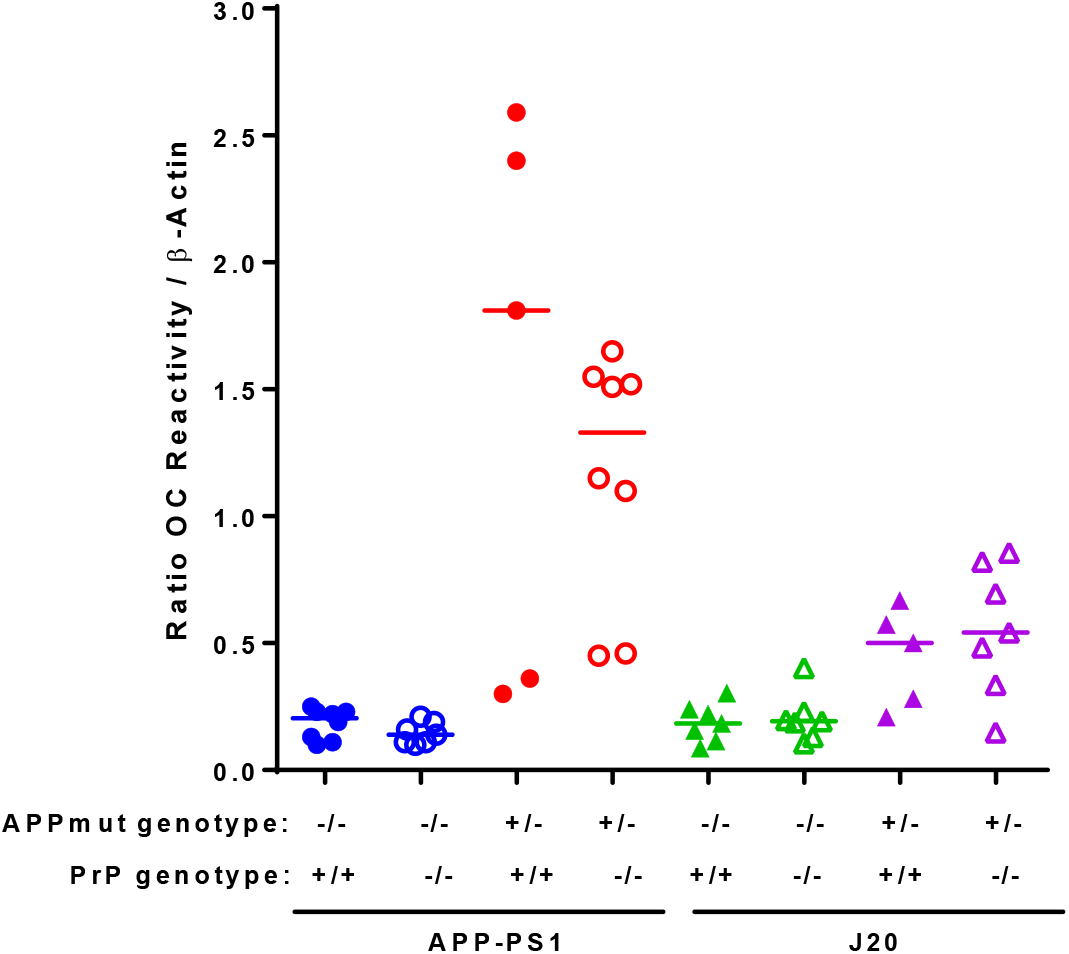
Aβ oligomers present in APP-PS1 and J20 mouse brain have different conformations, independent of PrP^C^ expression. Total brain homogenates were quantified by dotblot using OC antibody (WT vs APP-PS1, p=0.012; APP-PS1 vs APP-PS1 PrP^C^ KO, p>0.99; WT vs J20, p=0.87; J20 vs J20 PrP^C^ KO, p>0.99).

**Figure 8.**
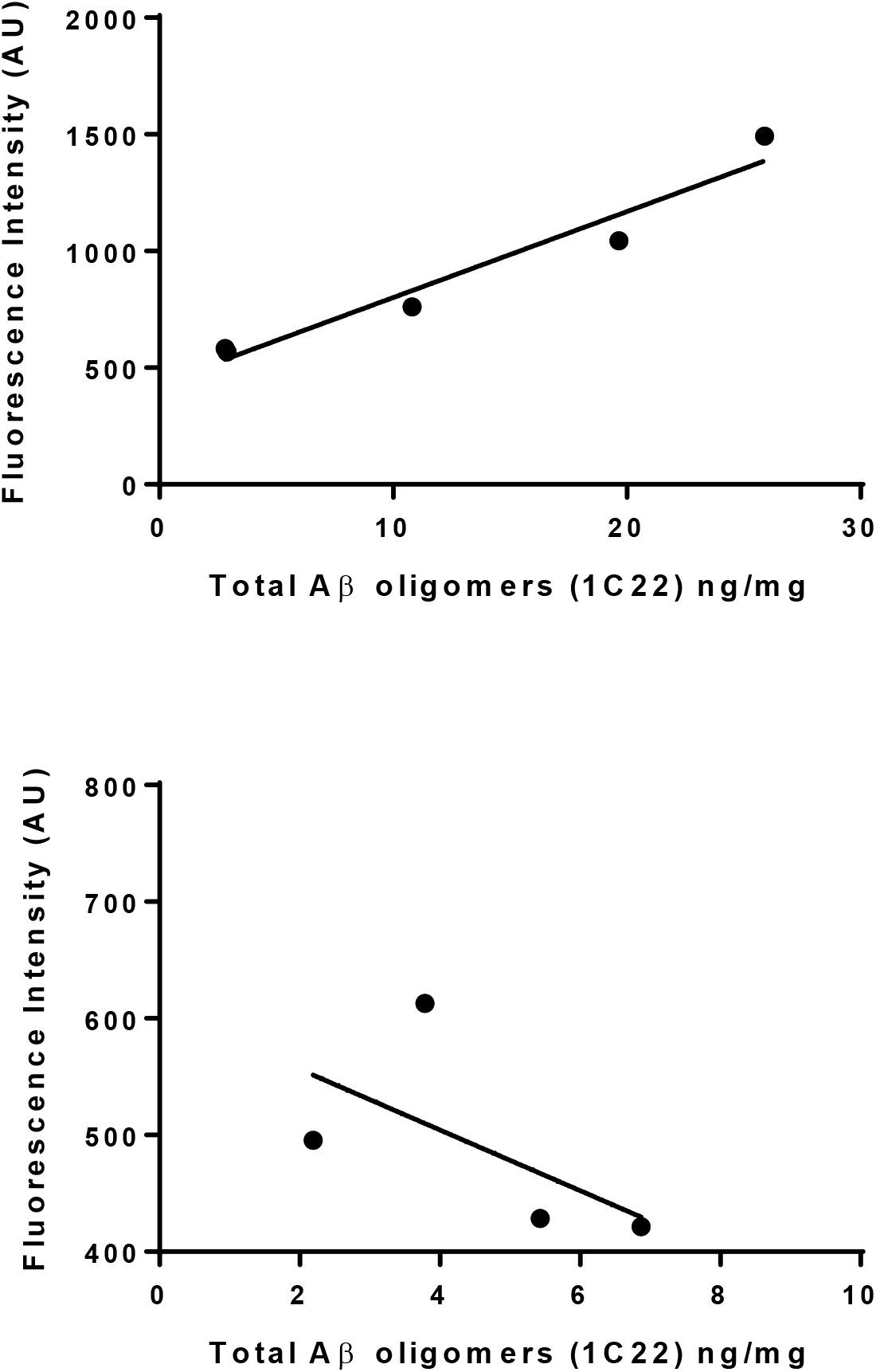
Positive correlation of Aβ oligomers that bind PrP^C^ with total amount of Aβ oligomers in APP-PS1, but not in J20 mice. **A)** Data from figures 5 and 6 showed a positive correlation for APP-PS1 brain samples (Pearson r=0.973, p=0.0052). **B)** No correlation was found for values obtained using J20 brain samples (Pearson r=-0.593, p=0.407).

We tested the ability of the four mouse lines to achieve several behavioural tasks at 3, 6 and 12 months of age. Healthy mice nest and burrow during the night time, when they are more active. All the mouse lines tested performed well at burrowing at all time points, except the J20 PrP^C^ KO mice, which exhibited an incremented disinterest on the task as age progressed when compared to the J20 mice (Figure 9). J20 PrP^C^ KO mice also showed an impairment on the Y-maze, a (hippocampal-dependent) short-term memory test (Figure 10). APP-PS1, APP-PS1 PrP^C^ KO, J20 mice and wild type mice performed equally well even after 12 months of age.

**Figure 9.**
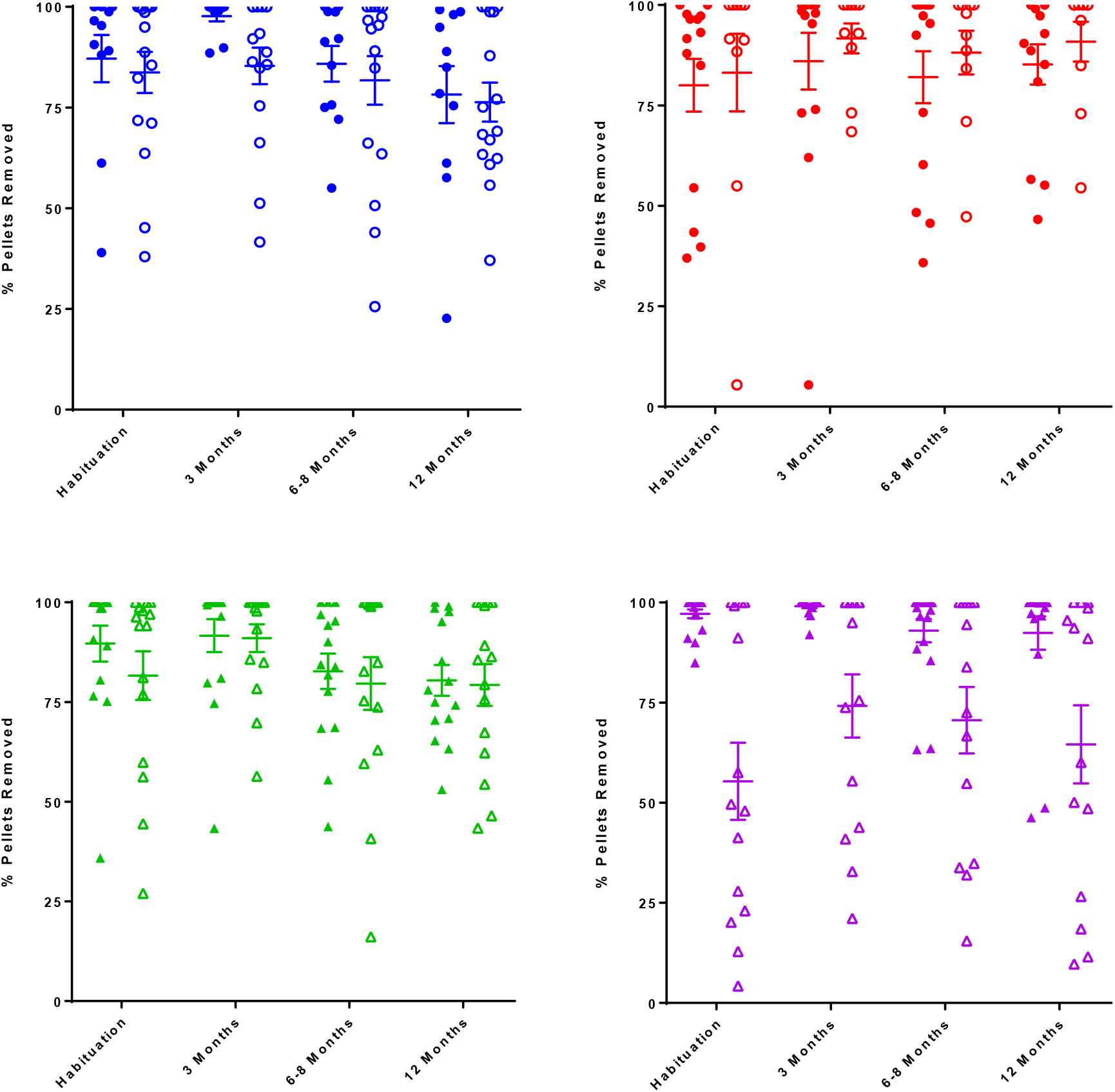
Ablation of PrP^C^ reduces burrowing activity in J20 mice after 3 months of age, but not in APP-PS1 model. Graph shows burrowing activity for **A)** WT and PrP^C^ KO mice (APP-PS1 littermates, blue), **B)** APP-PS1 and APP-PS1 PrP^C^ KO mice (red), **C)** WT and PrP^C^ KO (J20 littermates, green) and **D)** J20 and J20 PrP^C^ KO mice (purple) during the habituation phase and at 3, 6 and 12 months old. Significant differences were only found when comparing J20 to J20 PrP^C^ KO mice (at 3 months old: p=0.018; at 6 months old: p=0.04; at 12 months old: p=0.006). Data were analysed by two way ANOVA followed by Sidak’s post-hoc tests.

**Figure 10.**
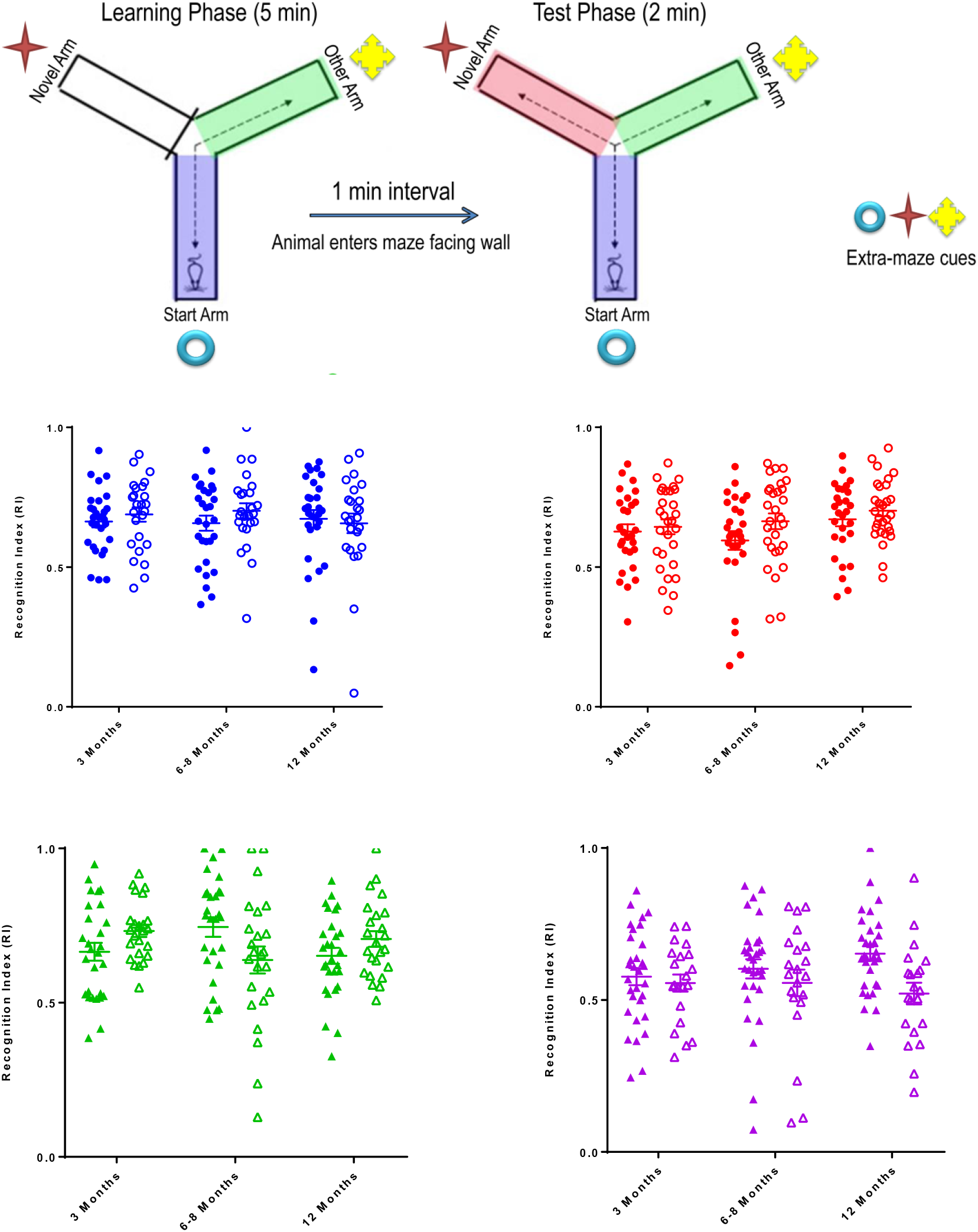
Lack of PrP^C^ affects performance of J20 mice in the Y-maze, but not that of APP-PS1 mice. **A)** Diagram depicts the Y-maze showing the geographical cues and methods for the test. Graph shows a novelty preference ratio obtained from mice **A)** WT and PrP^C^ KO mice (APP-PS1 littermates, blue), **B)** APP-PS1 and APP-PS1 PrP^C^ KO mice (red), **C)** WT and PrP^C^ KO (J20 littermates, green) and **D)** J20 and J20 PrP^C^ KO mice (purple) at 3, 6 and 12 months old when tested in the Y-maze mice. Significant differences were found when comparing J20 vs J20 PrP^C^ KO mice at 12 months old (p=0.013). Data were analysed by two way ANOVA followed by Sidak’s post-hoc tests.

On the novel object test, all mice performed well regardless of genotype showing a significant decline of novelty preference over time, indicating that this task was not sufficiently sensitive to discriminate between natural cognitive decline and AD related cognitive impairment (not shown).

Lastly, brain samples collected at 12 months were analysed using Golgi staining to provide an *in vivo* quantification of the dendritic spines present in the different mouse lines. Spines are postsynaptic structures, and a decreased spine density underlies cognitive deficits on behavioural tests (Spires-Jones and Knafo 2012). Pyramidal neurons on the CA1 hippocampal region presented a similar number of spines per μm when the basal dendrites were examined irrespective of either AD model or PrP^C^ expression, with the exception of a minor increase in spine density in J20 mice when compared with their wild-type littermates (WT vs J20 p=0.014) (Figure 11A, C). Similarly, a small increase in J20 apical spines was found when compared with the wild-type control (WT vs J20 p=0.038) and a partial reduction on spine density was evident on J20 PrP^C^ KO when compared with J20 mice (PrP^C^ KO vs J20 PrP^C^ KO, p=0.79; J20 vs J20 PrP^C^ KO, p=0.049) (Figure 11B, C).

**Figure 11.**
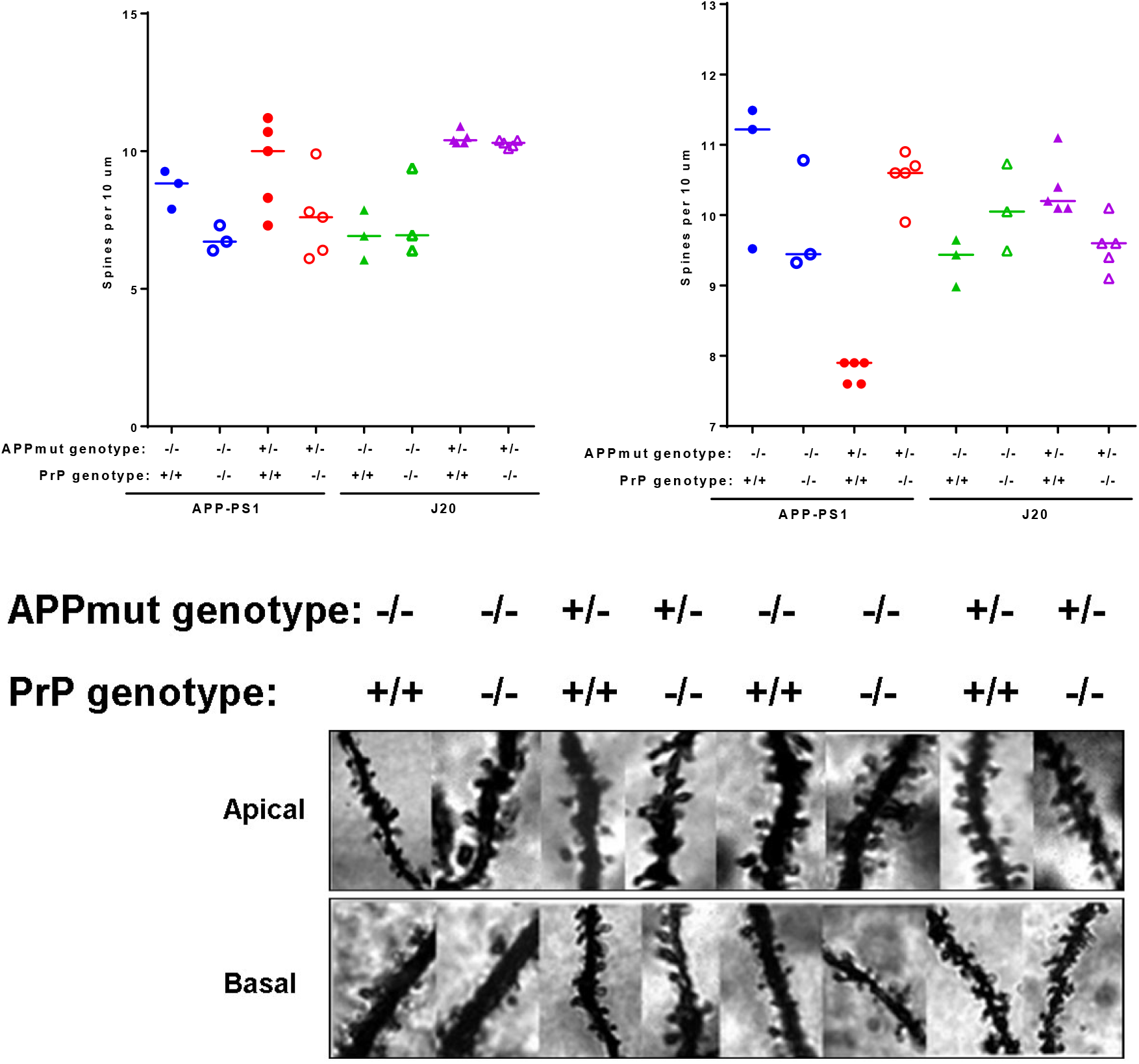
Effect of PrP^C^ expression on structural plasticity in both AD mouse models. **A)** Quantification of CA1 basal spines using Golgi staining (WT vs APP-PS1, p>0.99; PrP^C^ KO vs APP-PS1 PrP^C^ KO, p>0.99; WT vs J20 p=0.014; PrP^C^ KO vs J20 PrP^C^ KO, p=0.22; APP-PS1 vs APP-PS1 PrP^C^ KO, p=0.19; J20 vs J20 PrP^C^ KO, p>0.99). **B)** Quantification of CA1 apical spines using Golgi staining confirms that Aβ-induced spine loss is PrP-dependent on APP-PS1 model (WT vs APP-PS1, p=0.003; PrP^C^ KO vs APP-PS1 PrP^C^ KO, p=0.64; WT vs J20 p=0.038; PrP^C^ KO vs J20 PrP^C^ KO, p=0.79; APP-PS1 vs APP-PS1 PrP^C^ KO, p=0.001; J20 vs J20 PrP^C^ KO, p=0.049). **C)** Representative images of CA1 apical and basal dendritic spines of each group quantified.

A significant decrease in spine number was found on the apical dendrites of the APP-PS1 mice observed, versus wild-type controls (WT vs APP-PS1, p=0.003), suggestive of Aβ oligomer toxicity in dendrites in this model (Figure 11B, C). Interestingly, the ablation of PrP^C^ on this line rescued the spine loss on the apical dendrites, confirming that the Aβ-toxicity on dendritic spines is PrP^C^-dependent (PrP^C^ KO vs APP-PS1 PrP^C^ KO, p=0.64; APP-PS1 vs APP-PS1 PrP^C^ KO, p=0.001) (Figure 11B, C). Ablation of PrP^C^ in this model protected the spines from the soluble Aβ, confirming that the toxicity exercised by OC-conformational oligomers is mediated by PrP^C^.

## Discussion

Here, using a range of biochemical, histological and behavioural techniques we directly compared two different mouse models of AD: APP-PS1 and J20 mouse lines and the impact of PrP^C^ expression on them. We identified differences in the levels and localisation of Aβ plaques, peptides, oligomers, and PrP^C^-binding species between the two lines. However, PrP^C^ had no significant impact on these histological/biochemical parameters in either model. Interestingly, PrP^C^ deletion rescued spine loss in the APP-PS1 mouse but induced significant cognitive impairment in the J20 line, suggesting distinct interactions between Aβ and PrP^C^ could take place in these two mouse models. Further studies are needed to address this in more detail.

However, the observation that J20 mice have significantly lower quantity of Aβ oligomers and Aβ PrP^C^-binding species does provide a rationale for why Cisse *et al*. (Cisse et al. 2011) did not observe the PrP^C^-dependent phenotype reported in Gimbel *et al*. A biochemical study using several AD mouse models to compare the proportion of Aβ oligomers that interact with PrP^C^ from the total pool concluded that the amount of PrP^C^-binding Aβ varies from model to model. Unfortunately, the J20 model was not included in this study (Kostylev et al. 2015). Soluble fibrillar oligomers (OC-type) have been associated with increased levels of dementia and AD pathology (Tomic et al. 2009) and a study suggested that PrP^C^ has ∼30 times stronger affinity to OC-positive Aβ oligomers in comparison to A11-positive oligomers (Madhu and Mukhopadhyay 2020). All together, these results highlight the need to characterise Aβ oligomers present in the AD mouse models and AD brain samples to understand the mechanisms of Aβ toxicity in AD.

During Alzheimer’s disease there is a strong correlation between cognitive decline and loss of synapses. However, some AD mouse models fail to demonstrate this characteristic, with no correlation observed during quantification of dendritic spines and/or levels of synaptic proteins. Notwithstanding intensive efforts, the role of APP on structural spine plasticity is complex and not yet fully understood (Montagna, Dorostkar, and Herms 2017). A comparison of data obtained by several labs using the same Tg2576 mouse model, which overexpresses APP_Swe_ under the prion promoter, highlighted that the interpretation of the data is intricate and specification of age, gender, brain region and section of the dendrite quantified (apical or basal) are crucial to accurately unravel APP effects on spine plasticity (Jung and Herms 2012). In addition, some studies suggest that PS1 overexpression could lead to an elevated number of spines (Jung et al. 2011). As such, many studies have shown no changes or elevated number of spines in AD mouse models, with spatiotemporal patterns sometimes showing variability within the same study (Lanz, Carter, and Merchant 2003) making the results more difficult to interpret. In order to obtain an *in vivo* correlation of synaptic plasticity we analysed dendritic spines in the CA1 hippocampal region. Although this parameter has not been quantified previously in these mouse lines regarding the impact of PrP^C^ expression, *in vitro* experiments incubating wild-type mouse brain slices with Aβ oligomers indicated that Aβ-PrP^C^ binding induced loss of spines (Um et al. 2013; Um et al. 2012). Moreover, conditional deletion of *Prnp* after 12 and 16 months of age in APP-PS1 mice can reverse the behavioural deficits and recover the loss of synaptic proteins (Salazar et al. 2017). In the current study, a significant loss of spines on apical dendrites was found in APP-PS1 mice and was completely rescued by deletion of PrP^C^. These results confirm PrP-dependent Aβ toxicity in APP-PS1 mice. Surprisingly, the observed reduction in spines did not have an impact on the performance of behavioural tasks by the APP-PS1 mice, possibly due to the low difficulty of the challenges imposed. During behavioural tests, J20 PrP^C^ KO mice performed significantly worse than the J20 mice therefore we expected a reduction of dendritic spine density. However, the J20 mice showed a slight increase in the number of apical and basal spines, which was partially reduced by ablation of PrP^C^ only on the apical spines. As previously mentioned, an increase in spine density is sometimes found in AD mouse models possibly as a compensation mechanism. More detailed studies on innervated spine number, functional synapses and level of synaptic proteins could help to understand these results.

It is worth noting that the three behavioural tasks used differed from those used previously to show PrP^C^-dependent Aβ-toxicity (Gimbel et al. 2010). Unfortunately, we could not determine if ablation of PrP^C^ rescued the cognitive defects in APP-PS1 or J20 mice given that the tasks were possibly too simple, without sufficient sensitivity. We anticipate more challenging tests, such as the Morris water maze, would uncover the cognitive impairments present in these lines.

AD mouse models do not completely mimic the disease, whilst showing variable phenotypes between lines. Currently, the field is progressively moving towards knock-in models to avoid overexpression artefacts and potential confounds due to random genomic integration of transgenes. One such new knock-in model is the NL-F mouse line (Saito et al. 2014), which we are currently examining to understand physiological Aβ-PrP^C^ interactions.

In summary, our results agree with the published data on these two mouse lines and provide an explanation for previous contradictions, highlighting the need for thorough characterisation of Aβ oligomers, their binding to diverse receptors and subsequent effects on neurons. APP_swe_-PS1ΔE9 mice have proven to be an appropriate model to study Aβ-PrP^C^ interactions since they produce Aβ soluble fibrillary oligomers (OC-type) that bind PrP^C^, in contrast to J20 mice. As there are many and diverse Aβ aggregates present in the brain, identifying the toxic Aβ species in AD is paramount for selection of models that recapitulate specific conformers of interest.

## Acknowledgements

This work was funded by the UK Medical Research Council (MRC) and the Leonard Wolfson Experimental Neurology Centre. We thank staff of the MRC Prion Unit Biological Services facility for animal observation and care, to the Histology facility and M Ellis for technical assistance. We thank T Cunningham for reading the manuscript and helpful discussions.

## Conflict of interest

J.C. is a Director and shareholder of D-Gen Limited, an academic spin-out company working in the field of prion disease diagnosis, decontamination and therapeutics. D-Gen supplied antibody ICSM18.

## References

Benilova, I., E. Karran, and B. De Strooper. 2012. ‘The toxic Abeta oligomer and Alzheimer’s disease: an emperor in need of clothes’, Nat Neurosci, 15: 349–57.

Bozon, B., S Davis, and S. Laroche. 2003. ‘A requirement for the immediate early gene zif268 in reconsolidation of recognition memory after retrieval.’, Neuron, 40: 695–701.

Bueler, H., M. Fischer, Y. Lang, H. Bluethmann, H-P. Lipp, S.J. DeArmond, S.B. Prusiner, M. Aguet, and C. Weissmann. 1992. ‘Normal development and behaviour of mice lacking the neuronal cell-surface PrP protein’, Nature, 356: 577–82.

Cisse, M., P.E. Sanchez, D.H. Kim, K. Ho, G.Q. Yu, and L. Mucke. 2011. ‘Ablation of cellular prion protein does not ameliorate abnormal neural network activity or cognitive dysfunction in the j20 line of human amyloid precursor protein transgenic mice’, J Neurosci, 31: 10427–31.

Corbett, G. T., Z. Wang, W. Hong, M. Colom-Cadena, J. Rose, M. Liao, A. Asfaw, T. C. Hall, L. Ding, A. DeSousa, M. P. Frosch, J. Collinge, D. A. Harris, M. S. Perkinton, T. L. Spires-Jones, T. L. Young-Pearse, A. Billinton, and D. M. Walsh. 2020. ‘PrP is a central player in toxicity mediated by soluble aggregates of neurodegeneration-causing proteins’, Acta Neuropathol, 139: 503–26.

Cunningham, C., R. Deacon, H. Wells, D. Boche, S. Waters, C.P. Diniz, H. Scott, J.N. Rawlins, and V.H. Perry. 2003. ‘Synaptic changes characterize early behavioural signs in the ME7 model of murine prion disease’, Eur. J Neurosci, 17: 2147–55.

Der Laak, J.A., M.M. Pahlplatz, A.G. Hanselaar, and P.C. de Wilde. 2000. ‘Hue-saturation-density (HSD) model for stain recognition in digital images from transmitted light microscopy’, Cytometry, 39: 275–84.

Freir, D.B., A.J. Nicoll, I. Klyubin, S. Panico, J.M. Mc Donald, E. Risse, E.A. Asante, M.A. Farrow, R.B. Sessions, H.R. Saibil, A.R. Clarke, M.J. Rowan, D.M. Walsh, and J. Collinge. 2011. ‘Interaction between prion protein and toxic amyloid beta assemblies can be therapeutically targeted at multiple sites’, Nat Commun, 2: 336.

Gimbel, D.A., H.B. Nygaard, E.E. Coffey, E.C. Gunther, J. Lauren, Z.A. Gimbel, and S.M. Strittmatter. 2010. ‘Memory Impairment in Transgenic Alzheimer Mice Requires Cellular Prion Protein’, Journal of Neuroscience, 30: 6367–74.

Glabe, C.G. 2008. ‘Structural Classification of Toxic Amyloid Oligomers’, Journal of Biological Chemistry, 283: 29639–43.

Hu, N.W., A.J. Nicoll, D. Zhang, A.J. Mably, T. O’Malley, S.A. Purro, C. Terry, J. Collinge, D.M. Walsh, and M.J. Rowan. 2014. ‘mGlu5 receptors and cellular prion protein mediate amyloid-β-facilitated synaptic long-term depression in vivo’, Nat Commun, 5: 3374.

Jankowsky, J. L., and H. Zheng. 2017. ‘Practical considerations for choosing a mouse model of Alzheimer’s disease’, Mol Neurodegener, 12: 89.

Jankowsky, J.L., D.J. Fadale, J. Anderson, G.M. Xu, V. Gonzales, N.A. Jenkins, N.G. Copeland, M.K. Lee, L.H. Younkin, S.L. Wagner, S.G. Younkin, and D.R. Borchelt. 2004. ‘Mutant presenilins specifically elevate the levels of the 42 residue beta-amyloid peptide in vivo: evidence for augmentation of a 42-specific gamma secretase’, Human Molecular Genetics, 13: 159–70.

Jankowsky, J.L., A. Savonenko, G. Schilling, J. Wang, G. Xu, and D.R. Borchelt. 2002. ‘Transgenic mouse models of neurodegenerative disease: opportunities for therapeutic development’, Curr. Neurol. Neurosci. Rep, 2: 457–64.

Jarosz-Griffiths, H.H., E. Noble, J.V. Rushworth, and N.M. Hooper. 2015. ‘Amyloid-beta receptors: the good, the bad and the prion protein’, J Biol chem, 291: 3174–83.

Jung, C.K., M. Fuhrmann, K. Honarnejad, F. Van Leuven, and J. Herms. 2011. ‘Role of Presenilin1 in Structural Plasticity of Cortical Dendritic Spines in vivo’, J Neurochem, 119: 1064–73.

Jung, C.K., and J. Herms. 2012. ‘Role of APP for dendritic spine formation and stability’, Exp Brain Res, 217: 463–70.

Kayed, R., E. Head, F. Sarsoza, T. Saing, C.W. Cotman, M. Necula, L. Margol, J. Wu, L. Breydo, J.L. Thompson, S. Rasool, T. Gurlo, P. Butler, and C.G. Glabe. 2007. ‘Fibril specific, conformation dependent antibodies recognize a generic epitope common to amyloid fibrils and fibrillar oligomers that is absent in prefibrillar oligomers’, Molecular Neurodegeneration, 2.

Klyubin, I., A.J. Nicoll, A. Khalili-Shirazi, M. Farmer, S. Canning, A. Mably, J. Linehan, A. Brown, M. Wakeling, S. Brandner, D.M. Walsh, M.J. Rowan, and J. Collinge. 2014. ‘Peripheral Administration of a Humanized Anti-PrP Antibody Blocks Alzheimer’s Disease Abeta Synaptotoxicity’, J Neurosci, 34: 6140–45.

Koffie, R.M., M. Meyer-Luehmann, T. Hashimoto, K.W. Adams, M.L. Mielke, M. Garcia-Alloza, K.D. Micheva, S.J. Smith, M.L. Kim, V.M. Lee, B.T. Hyman, and T.L. Spires-Jones. 2009. ‘Oligomeric amyloid beta associates with postsynaptic densities and correlates with excitatory synapse loss near senile plaques’, Proc Natl Acad Sci U S A, 106: 4012–17.

Kostylev, M.A., A.C. Kaufman, H.B. Nygaard, P. Patel, L.T. Haas, E.C. Gunther, A. Vortmeyer, and S.M. Strittmatter. 2015. ‘Prion-Protein-Interacting Amyloid-beta Oligomers of High Molecular Weight are Tightly Correlated with Memory Impairment in Multiple Alzheimer Mouse Models’, J Biol. Chem, 290: 17415–38.

Lanz, T.A., D.B. Carter, and K.M. Merchant. 2003. ‘Dendritic spine loss in the hippocampus of young PDAPP and Tg2576 mice and its prevention by the ApoE2 genotype’, Neurobiol Dis, 13: 246–53.

Lauren, J., D.A. Gimbel, H.B. Nygaard, J.W. Gilbert, and S.M. Strittmatter. 2009. ‘Cellular prion protein mediates impairment of synaptic plasticity by amyloid-beta oligomers’, Nature, 457: 1128–32.

Liu, P., M.N. Reed, L.A. Kotilinek, M.K. Grant, C.L. Forster, W. Qiang, S.L. Shapiro, J.H. Reichl, A.C. Chiang, J.L. Jankowsky, C.M. Wilmot, J.P. Cleary, K.R. Zahs, and K.H. Ashe. 2015. ‘Quaternary Structure Defines a Large Class of Amyloid-beta Oligomers Neutralized by Sequestration’, Cell Rep, 11: 1760–71.

Madhu, P., and S. Mukhopadhyay. 2020. ‘Preferential Recruitment of Conformationally Distinct Amyloid-beta Oligomers by the Intrinsically Disordered Region of the Human Prion Protein’, ACS Chem Neurosci, 11: 86–98.

Montagna, E., M.M. Dorostkar, and J. Herms. 2017. ‘The Role of APP in Structural Spine Plasticity’, Front Mol Neurosci, 10: 136.

Mucke, L., E. Masliah, G.Q. Yu, M. Mallory, E.M. Rockenstein, G. Tatsuno, K. Hu, D. Kholodenko, K. Johnson-Wood, and L. McConlogue. 2000. ‘High-level neuronal expression of abeta 1-42 in wild-type human amyloid protein precursor transgenic mice: synaptotoxicity without plaque formation’, J Neurosci, 20: 4050–58.

Mullan, M., F. Crawford, K. Axelman, H. Houlden, L. Lilius, B. Winblad, and L. Lannfelt. 1992. ‘A pathogenic mutation for probable Alzheimer’s disease in the APP gene at the N-terminus of beta-amyloid’, Nat Genet, 1: 345–7.

Murrell, J., M. Farlow, B. Ghetti, and M. D. Benson. 1991. ‘A mutation in the amyloid precursor protein associated with hereditary Alzheimer’s disease’, Science, 254: 97–9.

Nicoll, A.J., S. Panico, D.B. Freir, D. Wright, C. Terry, E. Risse, C.E. Herron, T. O’Malley, J.D. Wadsworth, M.A. Farrow, D.M. Walsh, H.R. Saibil, and J. Collinge. 2013. ‘Amyloid-beta nanotubes are associated with prion protein-dependent synaptotoxicity’, Nat Commun, 4: 2416.

Purro, S. A., A. J. Nicoll, and J. Collinge. 2018. ‘Prion Protein as a Toxic Acceptor of Amyloid-beta Oligomers’, Biol Psychiatry, 83: 358–68.

Risse, E., A.J. Nicoll, W.A. Taylor, D. Wright, M. Badoni, X. Yang, M.A. Farrow, and J. Collinge. 2015. ‘Identification of a compound which disrupts binding of amyloid-beta to the prion protein using a novel fluorescence-based assay’, J Biol chem, 290: 17020–28.

Saito, T., Y. Matsuba, N. Mihira, J. Takano, P. Nilsson, S. Itohara, N. Iwata, and T.C. Saido. 2014. ‘Single App knock-in mouse models of Alzheimer’s disease’, Nat Neurosci, 17: 661–63.

Salazar, S.V., C. Gallardo, A.C. Kaufman, C.S. Herber, L.T. Haas, S. Robinson, J.C. Manson, M.K. Lee, and S.M. Strittmatter. 2017. ‘Conditional Deletion of Prnp Rescues Behavioral and Synaptic Deficits after Disease Onset in Transgenic Alzheimer’s Disease’, J Neurosci.

Sanderson, D.J., E. Hindley, E. Smeaton, N. Denny, A. Taylor, C. Barkus, R. Sprengel, P.H. Seeburg, and D.M. Bannerman. 2011. ‘Deletion of the GluA1 AMPA receptor subunit impairs recency-dependent object recognition memory’, Learn. Mem, 18: 181–90.

Smith, L. M., M. A. Kostylev, S. Lee, and S. M. Strittmatter. 2019. ‘Systematic and standardized comparison of reported amyloid-beta receptors for sufficiency, affinity, and Alzheimer’s disease relevance’, J Biol chem, 294: 6042–53.

Spires-Jones, T., and S. Knafo. 2012. ‘Spines, plasticity, and cognition in Alzheimer’s model mice’, Neural Plast, 2012: 319836.

Tomic, J.L., A. Pensalfini, E. Head, and C.G. Glabe. 2009. ‘Soluble fibrillar oligomer levels are elevated in Alzheimer’s disease brain and correlate with cognitive dysfunction’, Neurobiol Dis, 35: 352–58.

Um, J.W., A.C. Kaufman, M. Kostylev, J.K. Heiss, M. Stagi, H. Takahashi, M.E. Kerrisk, A. Vortmeyer, T. Wisniewski, A.J. Koleske, E.C. Gunther, H.B. Nygaard, and S.M. Strittmatter. 2013. ‘Metabotropic glutamate receptor 5 is a coreceptor for Alzheimer abeta oligomer bound to cellular prion protein’, Neuron, 79: 887–902.

Um, J.W., H.B. Nygaard, J.K. Heiss, M.A. Kostylev, M. Stagi, A. Vortmeyer, T. Wisniewski, E.C. Gunther, and S.M. Strittmatter. 2012. ‘Alzheimer amyloid-beta oligomer bound to postsynaptic prion protein activates Fyn to impair neurons’, Nat Neurosci, 15: 1227–35.

